## Supplementary material for "Wag31, a membrane tether, is crucial for lipid homeostasis in mycobacteria": supplimentary tables

**Table S1.** Plasmids and strains used in the study

| Plasmid | Description | Antibiotic resistance | Source |
|---|---|---|---|
| ***pJV53-Apra*** | pJV53 plasmid from Dr. Hatfull's lab wherein the *kan*$^r$ antibiotic cassette was swapped with apra$^r$ cassette. | apra | (1) |
| **pST-KiT$_{off}$** | L5 integrative pST-KiT plasmid harboring reverse tetR gene. | kan | (2) |
| **pST-gfp** | pST-KiT$_{off}$ construct harboring gfp between NdeI-HindIII sites. | Kan | This study |
| **pST-*wag31*** | pST-KiT$_{off}$ L5 integrative plasmid harboring FLAG-*wag31*-3X HA under the control of tetracycline induction | kan | This study |
| **pET-*wag31*** | pET28b harboring His-*wag31*-His | kan | This study |
| **pET-*wag31*$_{K20A}$** | pET28b harboring His-*wag31*$_{K20A}$-His | kan | This study |
| **pET-*wag31*$_{\Delta 1\text{-}60}$** | pET28b harboring His-*wag31*$_{\Delta 1\text{-}60}$-His | kan | This study |
| **pET-*wag31-gfp*** | pET28b harboring His-*wag31-gfp*-His | kan | This study |
| **pSCG-*wag31*** | Giles integrative plasmid harboring His-*wag31*-His under the control of wag31 native promoter | apra | This study |
| **pSCG- *wag31*$_{K20A}$** | Giles integrative plasmid harboring His-*wag31*$_{K20A}$-His under the control of wag31 native promoter | apra | This study |
| **pSCG- *wag31*$_{\Delta 1\text{-}60}$** | Giles integrative plasmid harboring His- *wag31*$_{\Delta 1\text{-}60}$-His under the control of wag31native promoter | apra | This study |
| **pNiT-*wag31*** | IVN inducible plasmid harboring 3X-FLAG-*wag31* | kan | This study |
| **pNiT-*murG*** | IVN inducible plasmid harboring 3X-FLAG-*murG* | kan | This study |
| **pNiT-*sepIVA*** | IVN inducible plasmid harboring 3X-FLAG- *sepIVA* | kan | This study |
| **pNiT-*msm2092*** | IVN inducible plasmid harboring 3X-FLAG- *msm2092* | kan | This study |
| **pNiT-*accA3*** | IVN inducible plasmid harboring 3X-FLAG- *accA3* | kan | This study |
| **pNiT-*mmpS5*** | IVN inducible plasmid harboring 3X-FLAG- *mmpS5* | kan | This study |
| **pNiT-*mmpL4*** | IVN inducible plasmid harboring 3X-FLAG- *mmpL4* | kan | This study |

| Strain | Description | Antibiotic resistance | Source |
|---|---|---|---|
| ***E. coli*** DH5α | *E. coli* strain used for cloning | none | Invitrogen |
| ***E. coli*** BL21DE3 | *E. coli* strain used for protein expression | kan | Novagen |
| ***Msm*** | *M. smegmatis* mc$^2$155 strain | none | ATCC |
| ***Msm::pJV53*** | *Msm* strain electroporated with pJV53 plasmid that expresses recJ and | apra | This study |
| ***Msm*::*gfp*** | *Msm* strain electroporated with pST-KiT-gfp | | |
| ***Msm*::pS-*wag31*** | *gp60* and *gp61* recombinase expressing Msm::pJV53 strain electroporated harboring integrative pST-KiT$_{off}$ | apra, kan | This study |
| **Δ*wag31*** | wag31 mutant in *Msm* in which the native *wag31* gene was replaced with hyg$^r$ marker. The strain | hyg, kan | This study |

| | | | |
|---|---|---|---|
| | harbours integrative tet inducible pST-*wag31* at L5 site. Cured to remove pJV53 episomal copy. | | |
| **Δ*wag31::wag31*** | Δ*wag31* electroporated with pSCG-wag31 | hyg, kan, apra | This study |
| **Δ*wag31::wag31*$_{K20A}$** | Δ*wag31* electroporated with pSCG-wag31$_{K20A}$ | hyg, kan, apra | This study |
| **Δ*wag31::wag31*$_{\Delta 1\text{-}60}$** | Δ*wag31* electroporated with pSCG-wag31$_{\Delta 1\text{-}60}$ | hyg, kan, apra | This study |
| ***Msm::wag31*** | *Msm* electroporated with pNiT-*wag31* | kan | This study |
| ***Msm::murG*** | *Msm* electroporated with pNiT-*murG* | kan | This study |
| ***Msm::sepIVA*** | *Msm* electroporated with pNiT-*sepIVA* | kan | This study |
| ***Msm::msm2092*** | *Msm* electroporated with pNiT-*msm2092* | kan | This study |
| ***Msm::accA3*** | *Msm* electroporated with pNiT-*accA3* | kan | This study |
| ***Msm::mmpS5*** | *Msm* electroporated with pNiT-*mmpS5* | kan | This study |
| ***Msm::mmpL4*** | *Msm* electroporated with pNiT-*mmpL4* | kan | This study |

**References**


1. J. C. van Kessel, G. F. Hatfull, Recombineering in Mycobacterium tuberculosis. *Nat Methods* **4**, 147-152 (2007).
2. V. Soni *et al.*, Depletion of M. tuberculosis GlmU from Infected Murine Lungs Effects the Clearance of the Pathogen. *PLoS Pathog* **11**, e1005235 (2015).

### Table S2: unique interacting partners found for Wag31

| Accession | Gene ID | # PSMs | # Unique Peptides | Localisation (Based on PMID: 34965429, 27114527, 20412585) | Functional category |
|---|---|---|---|---|---|
| A0QYB1 | MSMEG_3596 | 185 | 37 | Unknown | replication, recombination & repair |
| A0QTE7 | MSMEG_1813 | 104 | 16 | IMD | lipid metabolism |
| Q3L891 | MSMEG_0402 | 58 | 16 | PM-CW | cell wall & cell processes |
| A0R1Y2 | MSMEG_4915 | 61 | 16 | PM-CW | Intermediary metabolism and respiration |
| A0R152 | :MSMEG_4626 | 37 | 13 | IMD | Translation, ribosomal structure & biogenisis |
| A0QWG2 | MSMEG_2931 | 44 | 14 | PM-CW | Translation, ribosomal structure & biogenisis |
| A0QPS6 | MSMEG_0502 | 34 | 10 | unknown | Intermediary metabolism and respiration |
| A0R684 | MSMEG_6459 | 29 | 11 | Cytosolic | Intermediary metabolism and respiration |
| A0R582 | MSMEG_6099 | 41 | 10 | PM-CW | Cell wall and cell processes |
| A0R016 | MSMEG_4227 | 34 | 9 | IMD | Cell wall and cell processes |
| A0QQW2 | MSMEG_0897 | 20 | 7 | unknown | conserved hypothetical |
| A0QU64 | MSMEG_2092 | 41 | 7 | unknown | Cell wall and cell processes |
| A0R7C9 | MSMEG_6863 | 25 | 5 | unknown | cell wall & cell processes |
| A0QX15 | MSMEG_3138 | 28 | 4 | Cytosolic | Intermediary metabolism and respiration |
| A0QPE7 | MSMEG_0372 | 24 | 6 | IMD | Lipid metabolism |
| A0QV38 | MSMEG_2436 | 21 | 4 | IMD | unknown |
| A0QUZ5 | MSMEG_2393 | 24 | 6 | Cytosolic | Lipid metabolism |
| A0QV18 | MSMEG_2416 | 13 | 3 | Cell wall | cell wall & cell processes |
| A0QRA5 | EG_1046, MSMEG | 12 | 4 | Cell wall | Cell wall and cell processes |
| A0QUE0 | MSMEG_2174 | 25 | 7 | Cell wall | replication, recombination & repair |
| A0R2R8 | MSMEG_5211 | 23 | 6 | unknown | **Amino acid transport and metabolism** |
| A0R7J6 | MSMEG_6941 | 18 | 5 | IMD | conserved hypotheticals |
| A0QU53 | MSMEG_2080 | 18 | 6 | IMD | lipid metabolism |
| A0QQ17 | MSMEG_0594 | 16 | 6 | IMD | Intermediary metabolism and respiration |
| A0R3C8 | MSMEG_5427 | 20 | 7 | unknown | Intermediary metabolism and respiration |
| A0QVL0 | MSMEG_2611 | 18 | 4 | unknown | Intermediary metabolism and respiration |
| A0QXI0 | MSMEG_3309 | 10 | 3 | unknown | conserved hypothetical |
| A0QPG1 | MSMEG_0386 | 7 | 3 | unknown | Cell wall and cell processes |
| A0R4S6 | MSMEG_5937 | 18 | 4 | unknown | Intermediary metabolism and respiration |
| A0QV82 | MSMEG_2481 | 11 | 4 | unknown | conserved hypothetical |
| A0QWS9 | MSMEG_3051 | 16 | 3 | unknown | Intermediary metabolism and respiration |
| A0QWS2 | MSMEG_3044 | 14 | 4 | unknown | Intermediary metabolism and respiration |
| P0CH36 | MSMEG_1037 | 16 | 4 | unknown | Intermediary metabolism and respiration |
| A0R083 | MSMEG_4294 | 10 | 3 | unknown | Intermediary metabolism and respiration |
| A0QQX7 | MSMEG_0912 | 14 | 4 | unknown | lipid metabolism |
| A0QYE7 | MSMEG_3632 | 10 | 3 | PM-CW | Intermediary metabolism and respiration |
| A0QYG9 | MSMEG_3654 | 13 | 4 | PM-CW | cell wall & cell processes |
| A0QV36 | MSMEG_2434 | 10 | 3 | unknown | conserved hypotheticals |
| A0QVP7 | MSMEG_2648 | 11 | 4 | cell wall | lipid metabolism |
| A0QWX7 | MSMEG_3100 | 14 | 4 | unknown | Intermediary metabolism and respiration |
| Q3L887 | MSMEG_0406 | 16 | 3 | unknown | lipid metabolism |
| A0QX25 | MSMEG_3148 | 15 | 4 | Cell wall | conserved hypothetical |
| A0QZ46 | MSMEG_3894 | 10 | 3 | IMD | Intermediary metabolism and respiration |
| A0QUY3 | MSMEG_2379 | 10 | 3 | unknown | Intermediary metabolism and respiration |
| A0QRB9 | MSMEG_1061 | 10 | 3 | cell wall | replication, recombination & repair |
| A0R7D8 | MSMEG_6874 | 13 | 3 | unknown | Intermediary metabolism and respiration |
| Q9X5M1 | MSMEG_4958 | 16 | 4 | unknown | cell wall & cell processes |
| A0R7G3 | MSMEG_6901 | 14 | 3 | unknown | conserved hypothetical |
| A0R5K0 | MSMEG_6219 | 12 | 1 | PM-CW | regulatory protein |
| A0QW11 | MSMEG_2768 | 19 | 3 | unknown | anscription, replication, recombination & repair |
| A0R597 | MSMEG_6114 | 10 | 3 | unknown | Intermediary metabolism and respiration |
| A0QWN3 | MSMEG_3003 | 10 | 2 | unknown | Translation, ribosomal structure & biogenisis |
| A0QRD2 | MSMEG_1073 | 14 | 4 | IMD | Intermediary metabolism and respiration |
| A0R0N6 | MSMEG_4450 | 10 | 3 | IMD | replication, recombination & repair |
| A0QTF8 | MSMEG_1825 | 10 | 4 | unknown | Intermediary metabolism and respiration |
| A0R6C4 | MSMEG_6499 | 10 | 2 | unknown | conserved hypotheticals |
| A0QYQ7 | MSMEG_3746 | 12 | 3 | Cell wall | Intermediary metabolism and respiration |
| A0R184 | MSMEG_4659 | 12 | 3 | unknown | regulatory protein |
| A0R061 | MSMEG_4272 | 10 | 2 | cell wall | Intermediary metabolism and respiration |

| | | | | | |
|---|---|---|---|---|---|
| A0QXW7 | MSMEG_3451 | 10 | 2 | unknown | Intermediary metabolism and respiration |
| A0QYU0 | MSMEG_3785 | 9 | 3 | unknown | Unknown |
| A0R2Q7 | MSMEG_5199 | 9 | 2 | unknown | lipid metabolism |
| A0R1D1 | MSMEG_4709 | 8 | 3 | unknown | lipid metabolism |
| A0QTQ8 | MSMEG_1930 | 8 | 3 | IMD | replication, recombination & repair |
| A0R2Q5 | MSMEG_5197 | 8 | 3 | unknown | lipid metabolism |
| A0R3T9 | MSMEG_5592 | 8 | 2 | unknown | Intermediary metabolism and respiration |
| A0R6D9 | MSMEG_6514 | 8 | 2 | unknown | Virulence, detoxification, adaptation |
| A0QNZ7 | AMSMEG_0220 | 8 | 2 | IMD | Intermediary metabolism and respiration |
| A0QZ33 | MSMEG_3880 | 8 | 2 | unknown | Intermediary metabolism and respiration |
| A0R4P1 | MSMEG_5903 | 8 | 2 | unknown | Intermediary metabolism and respiration |
| A0QS40 | MSMEG_1340 | 8 | 2 | unknown | conserved hypotheticals |
| A0QQH7 | MSMEG_0759 | 8 | 2 | unknown | Intermediary metabolism and respiration |
| A0QSK8 | MSMEG_1514 | 8 | 2 | unknown | conserved hypotheticals |
| A0R6M2 | MSMEG_6598 | 8 | 2 | unknown | Intermediary metabolism and respiration |
| A0R436 | MSMEG_5691 | 7 | 3 | unknown | conserved hypothetical protein |
| A0R5X8 | MSMEG_6351 | 7 | 3 | unknown | Intermediary metabolism and respiration |
| A0QT17 | MSMEG_1679 | 7 | 2 | Cell wall | unknown |
| A0QXN2 | MSMEG_3363 | 7 | 2 | unknown | regulatory protein |
| A0R0W2 | MSMEG_4528 | 7 | 2 | unknown | Intermediary metabolism and respiration |
| A0R066 | MSMEG_4276 | 7 | 2 | unknown | Intermediary metabolism and respiration |
| A0QT08 | MSMEG_1670 | 6 | 3 | PM-CW | Intermediary metabolism and respiration |
| A0QPW2 | MSMEG_0539 | 6 | 3 | PM-CW | Intermediary metabolism and respiration |
| Q3I5Q7 | MSMEG_0919 | 6 | 3 | unknown | Cell wall and cell processes |
| A0QZ93 | MSMEG_3942 | 6 | 3 | unknown | conserved hypothetical protein |
| A0R6D7 | MSMEG_6512 | 6 | 2 | unknown | lipid metabolism |
| A0R071 | MSMEG_4282 | 6 | 2 | IMD | conserved hypothetical protein |
| A0R048 | MSMEG_4258 | 6 | 2 | PM-CW | Intermediary metabolism and respiration |
| A0R753 | MSMEG_6783 | 6 | 2 | unknown | Unknown |
| A0QRA8 | EG_1049;MSMEG | 6 | 2 | unknown | Conserved hypotheticals |
| Q9X5M0 | MSMEG_4959 | 6 | 2 | PM-CW | Translation, ribosomal structure & biogenisis |
| A0QUW2 | MSMEG_2357 | 6 | 2 | unknown | Intermediary metabolism and respiration |
| A0QNG7 | MSMEG_0035 | 6 | 2 | PM-CW | regulatory protein |
| A0QX83 | MSMEG_3207 | 6 | 2 | unknown | Intermediary metabolism and respiration |
| A0QVL6 | MSMEG_2617 | 6 | 2 | unknown | Intermediary metabolism and respiration |
| A0R170 | MSMEG_4645 | 6 | 2 | unknown | Intermediary metabolism and respiration |
| A0QYD4 | MSMEG_3619 | 6 | 2 | PM-CW | Intermediary metabolism and respiration |
| A0R2U7 | MSMEG_5239 | 6 | 2 | unknown | Intermediary metabolism and respiration |
| A0R6D6 | MSMEG_6511 | 6 | 2 | unknown | lipid metabolism |
| A0QWQ3 | MSMEG_3024 | 5 | 2 | unknown | Conserved hypotheticals |
| A0R177 | MSMEG_4652 | 5 | 2 | unknown | Unknown |
| A0QXB8 | MSMEG_3244 | 5 | 2 | unknown | conserved hypothetical protein |
| A0QTT2 | MSMEG_1954 | 4 | 2 | PM-CW | Cell wall and cell processes |
| A0QP17 | MSMEG_0240 | 4 | 2 | unknown | conserved hypothetical protein |
| A0QYT3 | MSMEG_3777 | 4 | 2 | unknown | Translation, ribosomal structure & biogenisis |
| A0R2L4 | MSMEG_5156 | 4 | 2 | unknown | replication, recombination & repair |
| A0R5I9 | MSMEG_6208 | 4 | 2 | unknown | lipid metabolism |
| A0QWU9 | MSMEG_3071 | 4 | 2 | PM-CW | Intermediary metabolism and respiration |
| A0QQL0 | MSMEG_0793 | 4 | 2 | unknown | Intermediary metabolism and respiration |
| A0QR20 | MSMEG_0956 | 4 | 2 | unknown | Intermediary metabolism and respiration |
| A0QVT2 | MSMEG_2685 | 4 | 2 | unknown | Conserved hypotheticals |
| A0R2Q6 | :MSMEG_5198 | 4 | 2 | unknown | lipid metabolism |
| A0QX95 | MSMEG_3219 | 4 | 2 | Cell wall | Intermediary metabolism and respiration |
| A0QP16 | MSMEG_0239 | 4 | 2 | unknown | Intermediary metabolism and respiration |
| A0QZY4 | MSMEG_4193 | 4 | 2 | Cytosolic | Conserved hypotheticals |
| A0R3L4 | MSMEG_5515 | 4 | 2 | Cytosolic | Intermediary metabolism and respiration |
| A0R5N7 | MSMEG_6256 | 3 | 2 | PM-CW | Intermediary metabolism and respiration |
